# Structural basis for flagellin induced NAIP5 activation

**DOI:** 10.1101/2023.06.13.544801

**Authors:** Bhaskar Paidimuddala, Jianhao Cao, Liman Zhang

**Affiliations:** Department of Chemical Physiology and Biochemistry, Oregon Health and Science University, Portland, OR 97239, USA

**Keywords:** NAIP/NLRC4 inflammasome, NAIP receptor, NLR proteins, Bacteria infection, Flagellin, cryo-EM

## Abstract

The NAIP/NLRC4 inflammasome is activated when NAIP binds to a gram-negative bacterial ligand. Initially, NAIP exists in an inactive state with a wide-open conformation. Upon ligand binding, the winged helix domain (WHD) of NAIP is activated and forms steric clash with NLRC4 to open it up. However, how ligand binding induces the conformational change of NAIP is less clear. To understand this process, we investigated the dynamics of the ligand binding region of inactive NAIP5 and solved the cryo-EM structure of NAIP5 in complex with its specific ligand, FliC from flagellin, at 2.93 Å resolution. The structure revealed a “trap and lock” mechanism in FliC recognition, whereby FliC-D0_C_ is first trapped by the hydrophobic pocket of NAIP5, then locked in the binding site by the insertion domain (ID) and C-terminal tail (CTT) of NAIP5. The FliC-D0_N_ domain further inserts into the loop of ID to stabilize the complex. According to this mechanism, FliC activates NAIP5 by bringing multiple flexible domains together, particularly the ID, HD2, and LRR domains, to form the active conformation and support the WHD loop in triggering NLRC4 activation.

## Introduction

The nucleotide-binding domain (NBD), leucine rich repeat (LRR) domain containing protein family (NLR family) apoptosis inhibitory proteins (NAIPs) constitute a family of cytosolic receptors that mediate host defense against Gram-negative bacteria(1). There are multiple NAIP proteins in mice to sense different bacterial ligands. For example, NAIP5 and NAIP6 are activated by flagellin, while NAIP2 and NAIP1 are activated by components of bacterial type III secretion system, the inner rod protein and needle protein, respectively(2, 3). Human genome only encodes one NAIP, which seems to be activated by all three ligands (4-6). Upon activation, NAIPs recruit and activate the NLR family CARD containing protein 4 (NLRC4). Structurally, both NAIP and NLRC4 contain the Nucleotide Binding Domain (NBD), Helical Domain 1 (HD1), Winged Helix Domain (WHD), Helical domain 2 (HD2), and Leucin Rich Repeat (LRR) domains. However, their N-terminal effector domains are different. NAIP has three Baculovirus IAP Repeat (BIR) domains, while NLRC4 has a Caspase Activation and Recruitment domain (CARD). Upon activation, NAIP recruits inactive NLRC4 and triggers its conformational change from the inactive state to the active state. Active NLRC4 further oligomerizes and nucleates the filamentation of caspase-1 (7-9), which eventually lead to the proteolytic activation of IL-1β, IL-18, GSDMD, and cause pyroptotic cell death(10-12).

NAIP and NLRC4 have highly similar active structures in their open conformation and form the NAIP/NLRC4 inflammasome complex(13, 14), but they must adopt a stable inactive conformation in steady cells to prevent auto-inflammation. NLRC4 achieves this with an auto-inhibited conformation that prevents it from oligomerizing. In a recent report, we presented the structure of inactive NAIP in a wide-open conformation (15). The distinct inactive conformations of NAIP and NLRC4 suggests that these two proteins must have different activation mechanisms, which leads to the question of how ligand binding induces the activation of NAIP. Previously, two cryo-EM structures containing NAIP5 bound to flagellin have been solved (13, 14), but the electron density in the ligand binding region was insufficient to support unambiguous model building and led to controversy in the interpretation of ligand-NAIP interaction (16).

To understand how ligand binding activates NAIP, we characterized the dynamics of the ligand-binding region of inactive NAIP5, and determined the high-resolution cryo-EM structure of NAIP5 in complex with FliC, the flagellin from *Salmonella typhimurium*, at 2.93 Å resolution. The resulting cryo-EM map allowed us to build atomic models around the ligand binding region with high confidence. By comparing the structures of unliganded and liganded NAIP5, we have provided molecular insights into how NAIP is activated by ligand binding.

## Results

### Flexibility of the C-terminal regions in unliganded NAIP5

We previously reported the cryo-EM structure of unliganded NAIP5 in a wide-open conformation(15). Unfortunately, despite achieving an overall resolution of 3.4 Å, the density of the HD2-LRR region was absent in the unliganded NAIP5 structure. Here we conducted further investigations to comprehensively understand the nature of this missing density. Using multi-body refinement in Relion(17), we characterized the dynamics of the HD2-LRR regions, and the result demonstrated that the missing regions in the HD2-LRR undergo approximately 30° movement (**Figure 1A, movie 1**). Considering that the dynamic regions in HD2-LRR primarily participate in ligand-binding, we suggest their flexibility is functionally important to ligand recognition.

**Figure 1.**
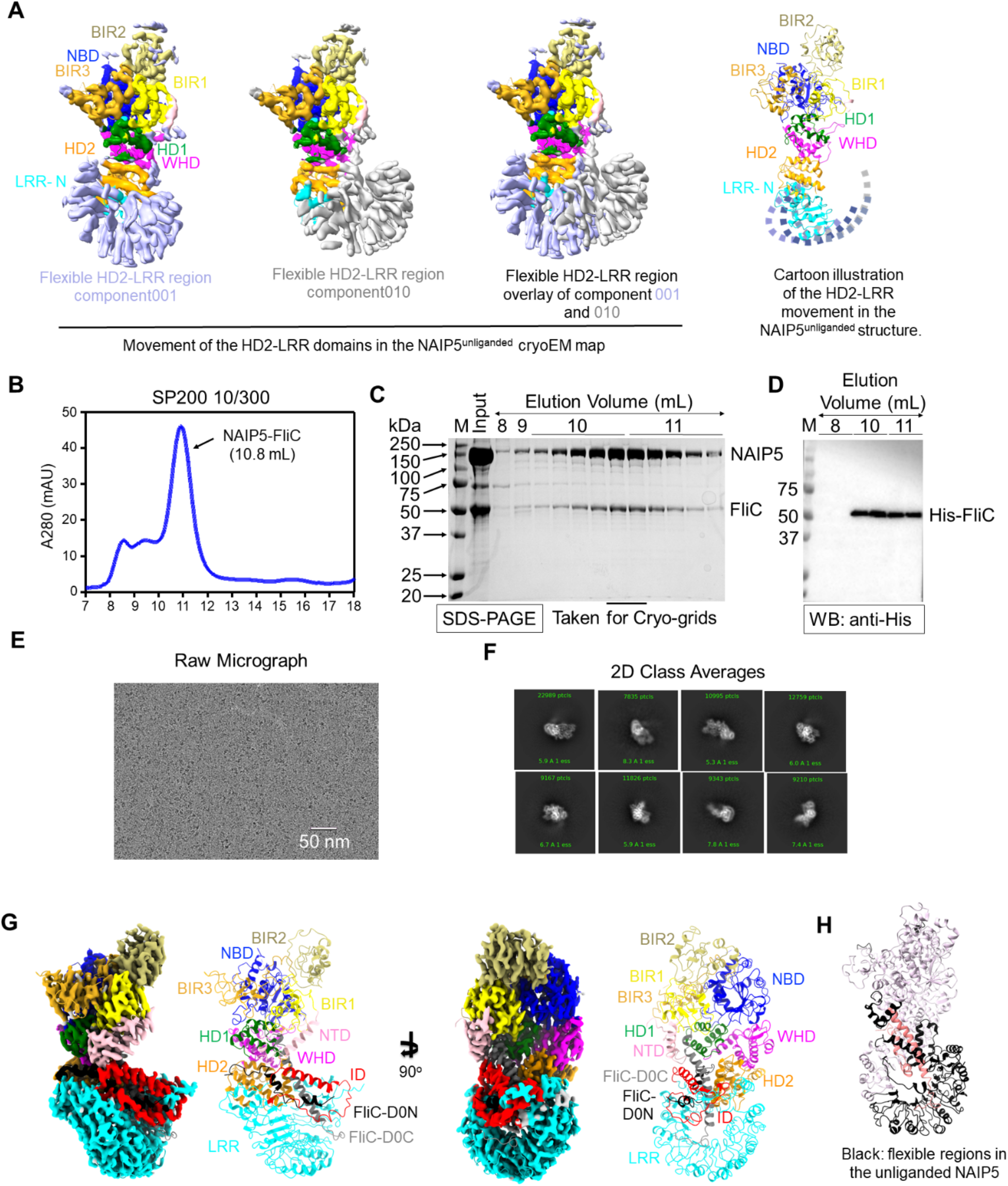
Dynamics of unliganded NAIP5, biochemical purification and cryo-EM structure of the NAIP5/FliC complex. A) Multi-Body Refinement of unliganded NAIP5 in Relion. B) The size exclusion chromatography profile of the NAIP5/FliC complex. C) SDS-PAGE of fractions in B. D) Western Blot of fractions in B. E) A representative cryo-EM micrograph and, F) 2D class averages showing side view and top/bottom views. G) Cryo-EM map and structure of the NAIP5/FliC complex, individual domains are color coded. H) The missing densities of unliganded NAIP5 (black) are plotted in the newly resolved NAIP5/FliC structure. FliC is shown in salmon.

We then attempted to analyze how ligand binding stabilizes the HD2-LRR regions by comparing our unliganded NAIP5 with two previous complexes of ligand/NAIP5/NLRC4 (referred to as PDB ID 6B5B(13) and 5YUD(14) hereafter). Unfortunately, the resolutions of ligand binding regions in both structures are insufficient for unambiguous model building. Consequently, the interpretations of the ligand-NAIP5 interaction in the two structures differ significantly (**Figure S1**) (13, 14). To support a mechanistic understanding of how the flexibility in HD2-LRR of NAIP5 accommodates ligand binding and how flagellin binding activates NAIP5, we decided to solve the structure of NAIP5/FliC complex at high resolution.

### Biochemical Purification and Cryo-EM studies of the NAIP5-FliC complex

Given that both 6B5B and 5YUD are solved in complex with NLRC4, which may induce sample heterogeneity and limit the high-resolution structure determination, we decided to take a different approach and reconstitute the complex with only NAIP5 and flagellin. Briefly, we co-expressed full-length NAIP5 with N-terminal Flag tag and the full-length *Salmonella typhimurium* flagellin, FliC with N-terminal His tag in Expi293F cells. Purification was carried out by anti-Flag affinity purification, followed by size exclusion chromatography (**Figure 1B)**. A large symmetric peak observed at 11 mL in the chromatogram of Superdex 200 showed the presence of NAIP5– FliC complex on the SDS-PAGE, which was further confirmed by western blotting (**Figure 1C,D)**. The corresponding peak fractions were taken for structure determination. Cryo-grids were screened and subjected to data collection under Titan Krios Transmission electron microscope equipped with a BioContinuum K3 direct electron detector.

Cryo-EM data processing was performed using cryoSPARC(18). Particles extracted from 4616 micrographs were subjected to multiple rounds of 2D classification before obtaining Ab-Initio maps. Next, the multiple rounds of heterogenous and homogenous refinement, followed by local refinement with a default mask resulted in a high-resolution map of 2.93 Å from the final particle set of 199,022 (**Figure 1E-G, S2)**.

To build the atomic model of active NAIP5, individual BIR domains, NBD, HD1 and WHD from the unliganded NAIP5 (PDB ID 7RAV), and HD2, LRR from the AlphaFold predicted structure (AF-Q9R016)(19) were fitted the into the cryo-EM map (15). FliC was built de-novo based on the side chain densities of cryo-EM map. The initial model of NAIP5/FliC complex was then subjected to multiple rounds of refinement using Coot (20) and Phenix real space refinement (21) (**Figure S3 A, B**).

The overall shape of our NAIP5/FliC structure is similar to the active NLRC4 (7, 8) and the NAIP5 subunits in both 6B5B and 5YUD **(Figure 1G, S1)**. To clarify one of the major differences between 6B5B and 5YUD, our structure showed residues 919-982 of NAIP5 forms insertion domain (ID) and directly contact with FliC. We also were able to confidentially build the ligand binding region with our high-resolution map, which is different in 6B5B and 5YUD, and mostly flexible in the unliganded NAIP5 structure (**Figure 1H**). More importantly, our structure reveals molecular details that allowed us to propose the mechanism underlying FliC induced NAIP5 activation.

### Structure of FliC in the NAIP5/FliC complex

Although we used full-length FliC in our sample preparation, the density map only revealed the D0_N_ helix and D0_C_ helix, which directly interacted with NAIP5. The remaining part of FliC appeared as a blob in the density map, likely because it is flexibly connected with D0_N_ and D0_C_ (**Fig. 2A**). In monomeric FliC, both D0_N_ and D0_C_ have been shown to be disordered (25), but they form a coiled-coil structure in the flagella filament (**Fig. 2B**) (22, 23). Our D0_N_ and D0_C_ structures in the NAIP5/FliC complex adopted a helical conformation but showed obvious differences from those in flagella, 6B5B and 5YUD (**Fig. 2B, Fig. S4 A, B**)(13, 14, 24). Rather than the coiled-coil structure in flagella, which is essential for the filament formation (22, 23), both D0_N_ and D0_C_ in the NAIP5/FliC complex bend in the middle and are separated by NAIP5 ID (**Fig. 2A**). Considering only the disordered structure in the monomeric D0_N_ and D0_C_ could allow them to separately bind NAIP5, these findings suggest NAIP5 may only recognize monomeric FliC.

**Figure 2.**
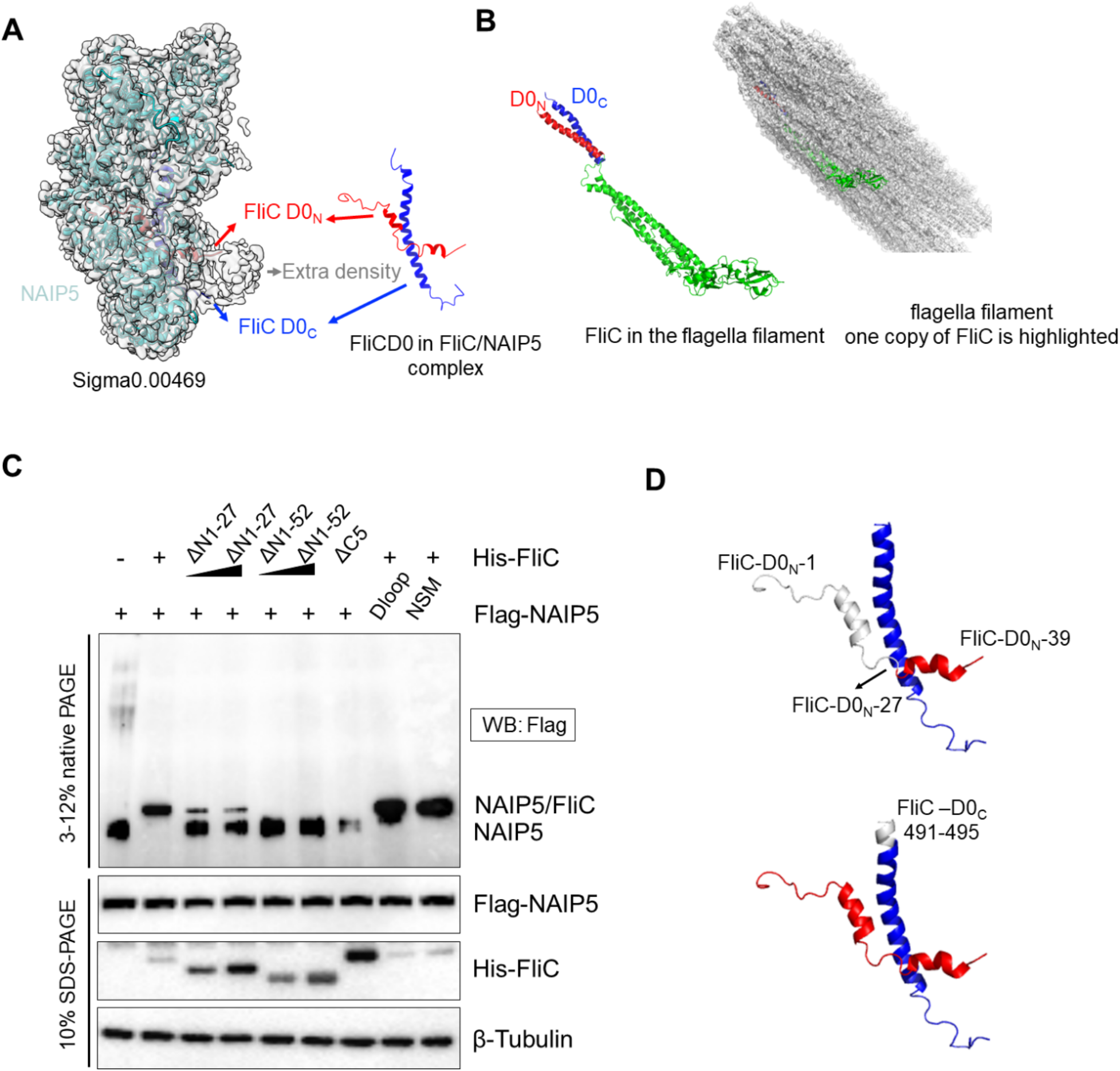
The FliC subunit in the NAIP5/FliC complex. A) The D0_N_ and D0_C_ helices of FliC in the cryo-EM map of NAIP5/FliC complex. The extra density that may correspond to part of remaining regions of FliC is labeled. B) D0_N_ and D0_C_ in flagella filament. C) The effect of different truncations of FliC on the complex formation. D) Illustration of the FliC truncations in C.

Interestingly, D0_N_ was considered dispensable in NAIP5 reorganization, as D0_C_ alone could activate the inflammasome to induce IL-1β maturation and cell death (13, 14). D0_N_ also does not directly interact with NAIP5 in 5YUD (**Fig. S4 B**). However, our structure showed extensive interactions between D0_N_ and NAIP5 ID, HD2 domains (**Fig. 2A, 3D**). We speculate that previous assays to measure the functional readout of the NAIP/NLRC4 inflammasome are less sensitive to define the interactions between NAIP5 and FliC, so we utilized Blue Native-PAGE (BN-PAGE) to directly monitor the formation of NAIP5/FliC complex, and tested the functional importance of D0_N_ and D0_C_ in NAIP5 binding. Compared with Full-length FliC, both deletion of

N-terminal 27 residues (DN27) or 52 residues (DN52) showed a greatly reduced super-shift band, which indicates D0_N_ /NAIP5 interaction is important for the complex formation (**Fig. 2C, D)**. D0_C_ is indeed critical, as deleting the 5 residues at the C-terminal end abolished complex formation (**Fig. 2C, D)**. The NAIP5-Dloop mutation and the nucleating surface mutation (NSM) that are unable to activate NLRC4(15), could form a normal complex with FliC, which agrees with their proposed functions in triggering NLRC4 conformational change (**Fig. 2C, D)**.

### Interactions between NAIP5 and FliC-D0_*C*_

In our structure, NAIP5 and FliC form extensive interactions, with both D0_N_ (residues 1-39) and D0_C_ (residues 453-495) embedded inside NAIP5 **(Fig. 1G, 2A)**. Consistent with 6B5B and 5YUD, the C-terminal residues (487-495) of D0_C_ insert into a hydrophobic pocket formed by the NTD, BIR1, and HD1 domains of NAIP5(**Fig 3A)**. This insertion is essential as deletion of the C-terminal 5 residues abolishes NAIP5/FliC interaction (**Fig 2C, 3D**). Notably, the central region of D0_C_ (470-486) is sandwiched by the helixes of ID and HD2 domains (**Fig 3B**), and the N-terminal end of D0_C_ (459-469) is packed by ID and LRR domains (**Fig 3C, S5**).

**Figure 3.**
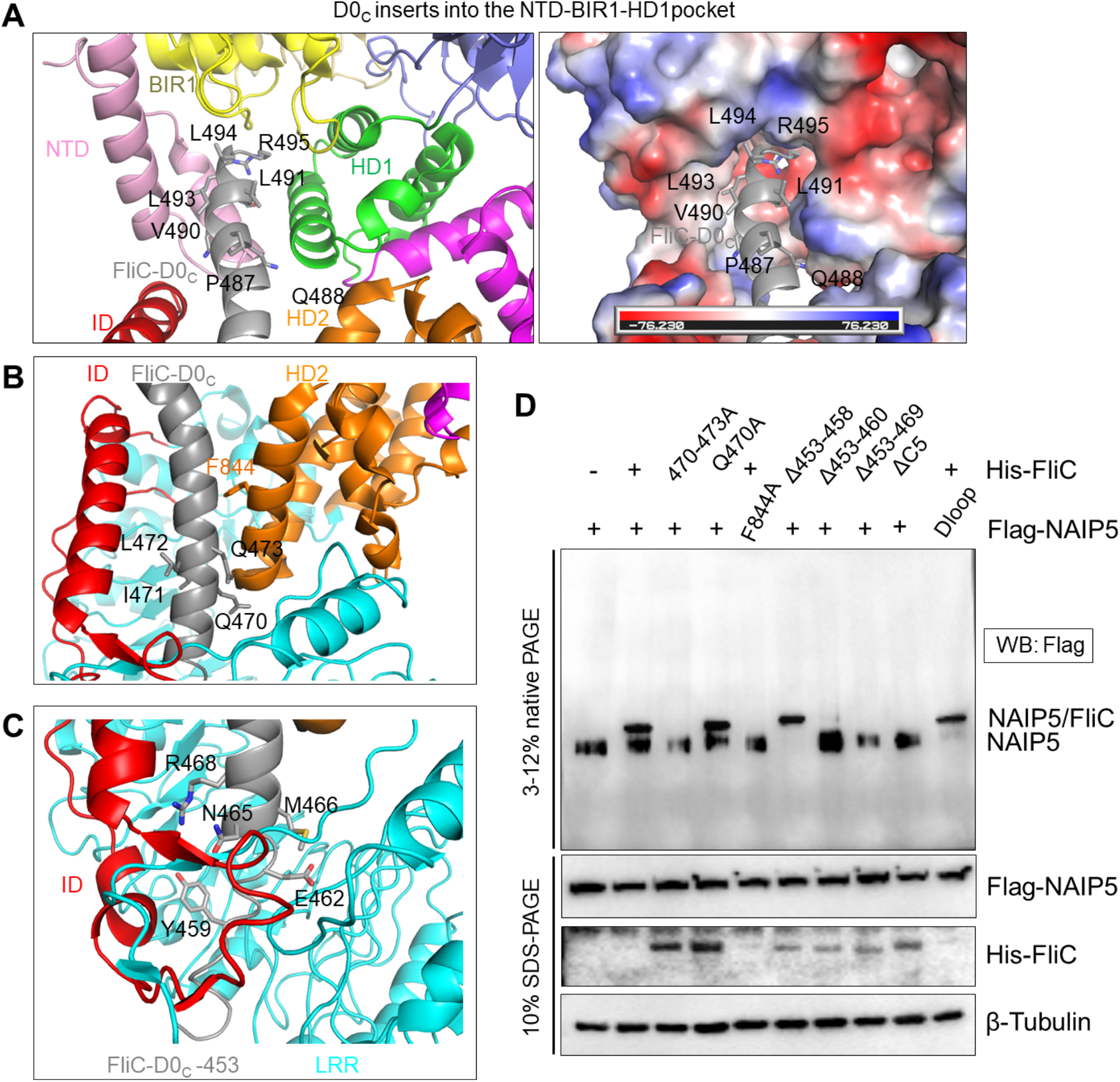
Interactions between FliC-D0_C_ and NAIP5. A) The C-terminal residues of D0_C_ insert into the hydrophobic pocket of NAIP5 formed by the NTD, BIR1, and HD1; NAIP5 domains are either color coded (left) or shown with their surface electrostatic potentials (right). D0_C_ is in grey. B) The middle part of D0_C_ (grey) is sandwiched by ID (red) and HD2 (orange). C) The N-terminal part of D0_C_ (grey) interacts with both ID (red) and LRR (cyan). D) The effect of different D0_C_ truncations in complex formation.

Except for the C-terminal residues, the importance of the middle part and N-terminal end of D0_C_ have not been extensively investigated. Thus, we designed mutations to test their ability to form complex with NAIP5. F844A in HD2 of NAIP5 was used as the positive control as this residue was previously mapped to be important for FliC-induced IL-1β processing (14). Indeed, F844A abolished complex formation on BN-PAGE (**Fig 3D**). Regarding FliC mutations, changing residues 470-473 into Alanine abolished complex formation (**Fig 3D**), suggesting the middle part of FliC is essential for NAIP5 recognition. At the N-terminal end, the removal of residues 453-458 has no effect, while the deletion of 453-460 greatly reduced, and the deletion of 453-469 completely abolished complex formation (**Fig 3D**), suggesting the packing of residues 459-469 by ID and LRR domains are functionally important.

### NAIP5-CTT inserts into NAIP5-ID loop to mediate FliC recognition

Having established the importance of the residues 459-469 of D0_C_ in FliC/NAIP5 recognition, we went on to analyze the molecular interactions in this region. It’s quite interesting that not only D0_C_ simultaneously bind ID and LRR, but the C-terminal tail (CTT) of LRR also inserts into the ID loop (**Fig 4A**), which is a feature that has not been overserved in other NLRs, to the best of our knowledge. Deletion of the CTT residues from 1388 to 1403 abolishes the NAIP5/FliC complex formation. Among them, residues 1388-1398 that interact with ID and FliC may be more important, as Δ1398-1403 can still bind FliC, although at a reduced level (**Fig 4B**). To further narrow down key residues in NAIP5-LRR and CTT, we designed more mutations on NAIP5, including L1393, I1394 to ED (LI-ED); M1361, L1362 to ED (ML-ED); L1365 to A; I1388, I1389, F1390 to DDH (IIF-DDH); K1392, L1393, I1394 to AGS (KLI-AGS). All the mutations abolished the complex formation (**Fig 4A, B**).

**Figure 4.**
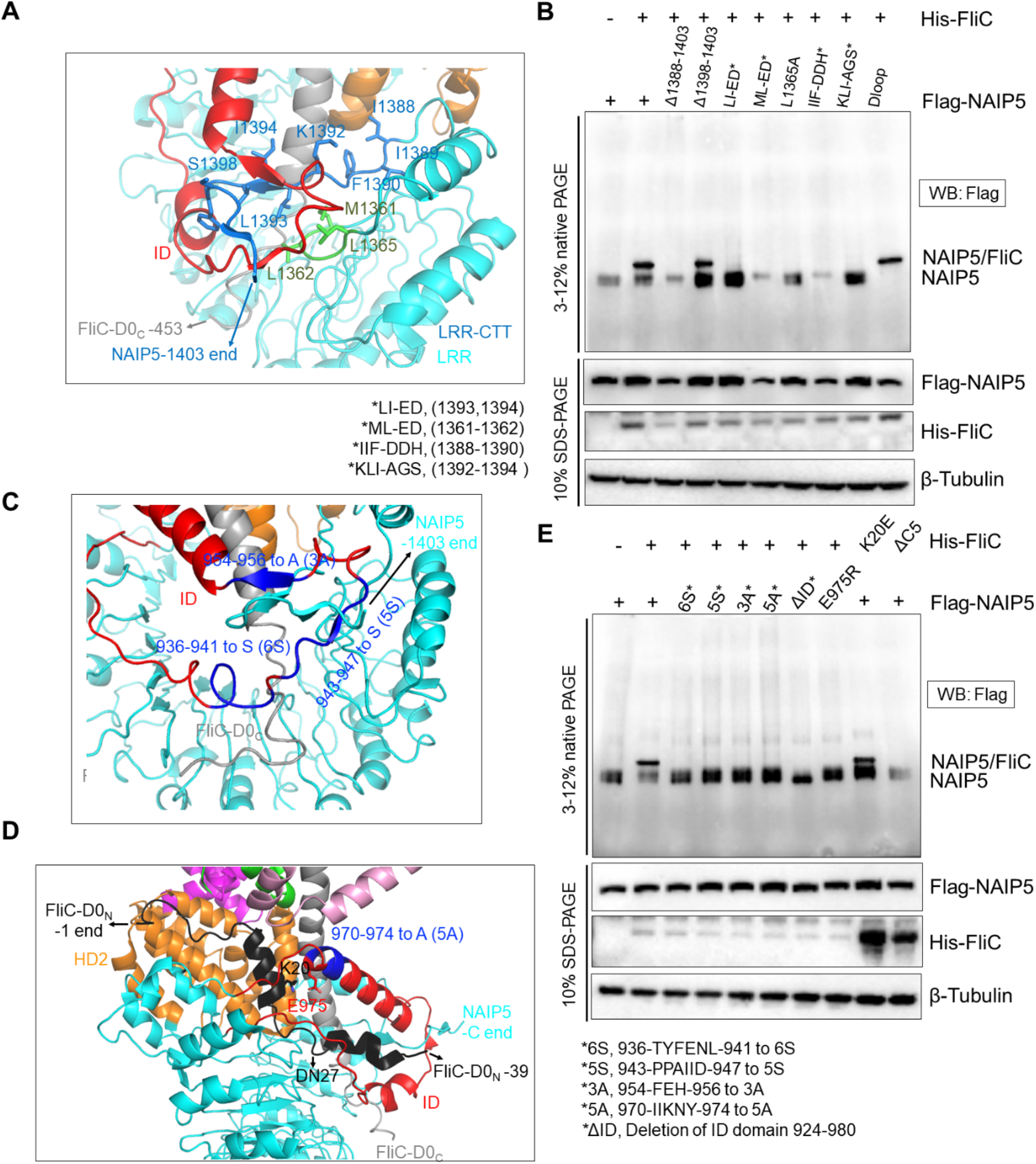
Interactions in the ID/CTT/D0_C_ regions. A) the CTT of NAIP5 inserts into the ID loop. B) Effects of mutations in the CTT in the complex formation. C) Residues mapped in the ID loop that are involved in ID/CTT and ID/D0_C_ interactions. D) D0_N_ inserts into ID loop and interacts with both ID and HD2. E) Effects of mutations in C and D in the complex formation.

Next, we mapped the key residues in ID region by separately mutating the ID fragments. Specifically, we changed 936-TYFENL-941 to 6S (6S), 943-PPAIID-947 to 5S (5S), and 954-FEH-956 to 3A (3A). All of them abolished the complex formation (**Fig 4C, E**). Altogether, these data suggest that NAIP5/FliC interaction is highly sensitive to the change of sequences in the ID and CTT regions, indicating an extensive network of ID-D0_C_-CTT interactions together determines the stability of NAIP5/FliC complex.

### Interactions between NAIP5 and FliC-D0_N_

In our structure, D0_N_ is inserted into the ID domain and makes contact with both the ID and HD2 domains (**Fig 4D)**. We have shown before that the DN27 truncation of FliC has a reduced ability to bind to NAIP5 (**Fig. 2C, D**). Consistently, the K20 on D0_N_ form charge-charge interaction with E975 of NAIP5, and K20E reduced but not abolished FliC/NAIP5 interaction. In comparison, mutations on NAIP5 residues caused a more severe phenotype. For example, both E975R and 970-IIKNY-974 to 5A (5A) mutations that may disrupt D0_N_-ID interaction abolished complex formation (**Fig 4D, E)**.

### Proposed mechanism for flagellin induced NAIP5 activation

With these analyses, we propose the mechanism for flagellin-induced NAIP5 activation. Overall, the recognition of FliC by NAIP5 involves three steps: D0_C_ Trap, CTT-ID lock, and D0_N_ guard.

Because the hydrophobic pocket is the only FliC binding element that pre-exists in the inactive NAIP5, the hydrophobic pocket and D0_C_ interaction is likely the first step in the recognition process. Once the C-terminal residues of D0_C_ are inserted into the hydrophobic pocket, the remaining portion of it forms multiple interactions with the flexible HD2, ID, and LRR domains to create a more rigid structure. This brings the CTT and ID of NAIP5 closer to each other, allowing CTT to insert into the ID loop and lock the D0_C_ insertion channel, keeping NAIP in the active conformation. Finally, D0_N_ inserts into ID, further stabilizing the complex. Altogether, the binding of FliC brings the HD2 and LRR domains of NAIP5 to their active positions, indirectly induces the movement of WHD, and forces the 17-18 loop to clash with NLRC4 to open it up (**Fig. 4, movie 2**), and eventually initiate the inflammasome signal pathway.

Surprisingly, although the unliganded NAIP5 structure showed a clear ATP in the nucleotide-binding pocket (15), we did not observe any nucleotide density in the same region of the active NAIP5/FliC complex. Consistently, the conformation of Walker A motif (470-**G**ETGS**GKT**-477) seems more flexible in the NAIP5/FliC complex, and appears different between these two maps (**Figure S5**), which suggest the ATP is hydrolyzed or released in our sample upon ligand binding. However, previous study has shown that mutating the Lysine residue in the walker A motif to Arginine, which reduces ATP binding, does not abolish the activation of NAIP/NLRC4 pathway(25). The ATPase activity of NAIP5 is moderate and is not enhanced by FliC (15). What’s more, a reliable nucleotide density was observed in the NAIP5/FliC complex of 5YUD(14), indicating that neither ATP hydrolysis nor release is essential for NAIP5 activation **(Figure S5)**. The differences in ATP binding in our sample and 5YUD could potentially be caused by different expression systems. Taken together, these findings suggest that the function of nucleotide binding in NAIP may not be as crucial as it is in other NLRs (26, 27).

## Discussion

In this study, we proposed the mechanism of FliC-induced NAIP5 activation. Our high-resolution cryo-EM structure of the NAIP5/FliC complex provides detailed insights into the recognition of flagellin by NAIP5, and the subsequent activation of NAIP5. Interestingly, both the D0_N_ and D0_C_ regions of FliC and the ligand-binding regions of NAIP5 exhibit flexibility before their interaction. Structural rearrangements occur in both components during the recognition process, suggesting that the initial FliC recognition may involve weak interactions that are strengthened through structural rearrangement. Additionally, the flexibility of the ligand-binding region in unliganded

NAIP5 provides the space for initial contact with FliC, enabling it to be trapped in the hydrophobic pocket within the NTD-BIR1-HD1 domains. Furthermore, the insertion of NAIP5-CTT into the ID loop locked D0_c_ into its binding pocket, and interactions between the ID and D0_N_ likely contribute to an additional stabilization mechanism. Overall, these interactions directly induce the position of the ID, HD2, and LRR domains at their active conformation, leading to a steric clash between WHD and NLRC4.

It is quite interesting to observe the incorporation of CTT into the ID loop following the binding of the ligand. LRR is commonly used as a ligand binding domain in Toll-like receptors (TLRs) and NLRs. But this is the first time the CTT was observed to participate in ligand binding. More importantly, a functional CTT is essential for stabilizing the NLR proteins and the NAIP/ligand complex. Among the currently solved mammalian NLR structures, NLRP9 also has a C-terminal tail and interacts with its LRR(28). It will be interesting to see if the tail appears in other NLRs and if they are also involved in ligand binding.

## Acknowledgments

We thank Claudia Lopez and Steven Adamou at the Multiscale Microscopy Core of OHSU; Janette Myers, Sean Mulligan, Craig Yoshioka, and Vamseedhar Rayaprolu at the Pacific Northwest Center for Cryo-EM (PNCC) for their help in cryo-electron microscopy operation. A portion of this research was supported by NIH grant U24GM129547 and performed at the PNCC at OHSU and accessed through EMSL (grid.436923.9), a DOE Office of Science User Facility sponsored by the Office of Biological and Environmental Research. This work was supported by the National Institutes of Health grant R01AI165580 (L.Z.). We apologize to authors whose work could not be cited because of space limitation.

## Data availability

The cryo-EM map of the protein complex was deposited in the Electron Microscopy Data Bank under the accession ID: EMD-29296 and the atomic coordinates were deposited in the PDB under the accession ID: 8FML.

## Author contributions

B.P., L.Z. and J.C. conceived the study, designed the experiments and analyzed the data. B.P. reconstituted the protein complex, performed cryo-EM experiments and refined the structure.

B.P. and L.Z. analyzed the structure and designed structure-based mutants. B.P. performed biochemical assays. J.C. analyzed the dynamics of unliganded NAIP5. B.P., L.Z. and J.C. wrote the manuscript.

